# Encoding of 3D Head Orienting Movements in Primary Visual Cortex

**DOI:** 10.1101/2020.01.16.909473

**Authors:** Grigori Guitchounts, Javier Masis, Steffen BE Wolff, David Cox

## Abstract

Animals actively sample from the sensory world by generating complex patterns of movement that evolve in three dimensions. At least some of these movements have been shown to influence neural codes in sensory areas. For example, in primary visual cortex (V1), locomotion-related neural activity influences sensory gain, encodes running speed, and predicts the direction of visual flow. As most experiments exploring movement-related modulation of V1 have been performed in head-fixed animals, it remains unclear whether or how the naturalistic movements used to interact with sensory stimuli– like head orienting–influence visual processing. Here we show that 3D head orienting movements modulate V1 neuronal activity in a direction-specific manner that also depends on the presence or absence of light. We identify two largely independent populations of movement-direction-tuned neurons that support this modulation, one of which is direction-tuned in the dark and the other in the light. Finally, we demonstrate that V1 gains access to a motor efference copy related to orientation from secondary motor cortex, which has been shown to control head orienting movements. These results suggest a mechanism through which sensory signals generated by purposeful movement can be distinguished from those arising in the outside world, and reveal a pervasive role of 3D movement in shaping sensory cortical dynamics.

## Introduction

Animals sample the visual world by producing complex patterns of movements^1^. Primates, birds, cats, fish, and insects move their eyes, heads, or bodies to stabilize the gaze, shift gaze, sample the visual scene, or estimate distance using parallax motion cues^2–6^. Saccadic eye movements in primates are accompanied by a suppression of neural activity in visual areas, which serves to reduce blur during fast movement and produce an active sampling of the scene^6–14^. Freely-moving rodents also make complex patterns of head and eye movements^4,5,15–19^. For example, rodents use head movements to estimate distances when jumping across a gap^3,20^, and rats move their eyes to stabilize their gaze^17,18^. Similarly, mice also move their eyes under head-fixed conditions^21,22^. However, the neural consequences of many of these movements remain unclear.

Despite the myriad of movements animals make to support visual perception, vision has typically been studied by presenting stimuli to restrained animals. In this framework, vision is a passive sense and visual processing is a feedforward computation in which information enters the brain through the retina and is processed one stage at a time as it travels deeper into the brain, through thalamus and cortex^23–26^. Consistent with this view, the majority of neurophysiological experiments on primary sensory cortical areas have found that information in those areas is chiefly sensory in nature.

Importantly, visual cortical areas receive a multitude of feedback and modulatory inputs that influence sensory processing; this feedback may afford visual areas with information required to distinguish between self-generated sensory signals and those that arise in the outside world^27–29^. One candidate origin for efference copy signals in visual cortex is secondary motor cortex (M2), a medial prefrontal structure linking multiple motor and sensory regions^30^. Anatomical evidence has suggested that M2 is a motor associational region because it projects both to subcortical targets such as superior colliculus^30–32^, brainstem nuclei controlling eye movements^33,34^, and the spinal cord^30,35,36^, as well as multimodal cortical areas such as primary sensory, retrosplenial, and posterior parietal cortices^30^. Furthermore, M2 has extensive bidirectional connections with visual cortex^30,32,37^. Functional evidence further supports the idea of M2’s involvement in orienting movements. Electrical stimulation of M2 results in eye and orienting movements^22,38,39^, usually contralateral to the stimulated hemisphere, while optogenetic stimulation of M2 axons over V1 leads to turning behavior in head-fixed mice^40^. Conversely, temporary unilateral inactivation of M2 leads to contralateral neglect in a memory-guided orienting task^41^, while unilateral lesions of M2 result in hemilateral neglect^42^. Neuronal activity correlates of orienting movements have also been recorded in M2, both in the context of behavioral tasks^22,41^ and in freely orienting animals^43,44^. These anatomical, physiological, and functional lines of evidence implicate M2 as a potential source of movement-related signals in V1, which may be used to process self-generated sensory signals.

One defined means through which sensory cortices encode movement signals is via a unidimensional state variable that corresponds to the animal’s overall locomotion. Such movement-related signals have been found in a myriad of sensory brain regions^19,27,40,45–56^. This locomotion signal can have varied effects on sensory cortices. For example, while movement suppresses activity in mouse auditory cortex^27^, during locomotion on a treadmill cells in primary visual cortex (V1) increase their activity, reflect running speed, and increase the gain on encoding of visual stimuli^45–49^.

In addition to locomotion *per se* influencing neural activity, specific movements have been shown to modulate sensory cortical activity, including orofacial movements and movements associated with different directions of visual flow^40,54–56^. However, such studies have primarily been done in head-fixed rodents, which has made it difficult to answer whether sensory areas are modulated by naturalistic movements that support sensory sampling. By recording activity in V1 during free behavior in rats, here we demonstrate that head orienting movements cause suppression of neural activity in visual cortical neurons. Furthermore, V1 activity reflected the direction of head movement, both in the dark and in the light, with individual neurons tuned to the direction of movement. Responses to visual stimulation were reduced during orienting movements compared to rest, and encoding of movement direction depended on M2. Our results demonstrate that cortical sensory dynamics encode specific movements, and raise the possibility that head orienting movements in rodents serve the same functional purpose as saccades do in primates.

## Results

### Bidirectional Responses in the Light and Dark in V1 Encode 3D Head Orienting Movement Direction

To investigate the impact of spontaneous movements on sensory cortical dynamics, we recorded neuronal activity using tetrode arrays targeting layer 2/3 of rat V1 while the animals behaved freely in a home-cage arena (Figs. 1a-c, S1a-f). Movements were captured using a head-mounted inertial measurement unit (IMU) (see Materials and Methods). We focused our analysis on 3D orienting movements of the head, which we refer to as head orienting movements (HOMs), as well as total acceleration, which we term overall movement; this enabled us to distinguish general movement-related activity from activity related to head-based visual sampling.

**Figure 1:**
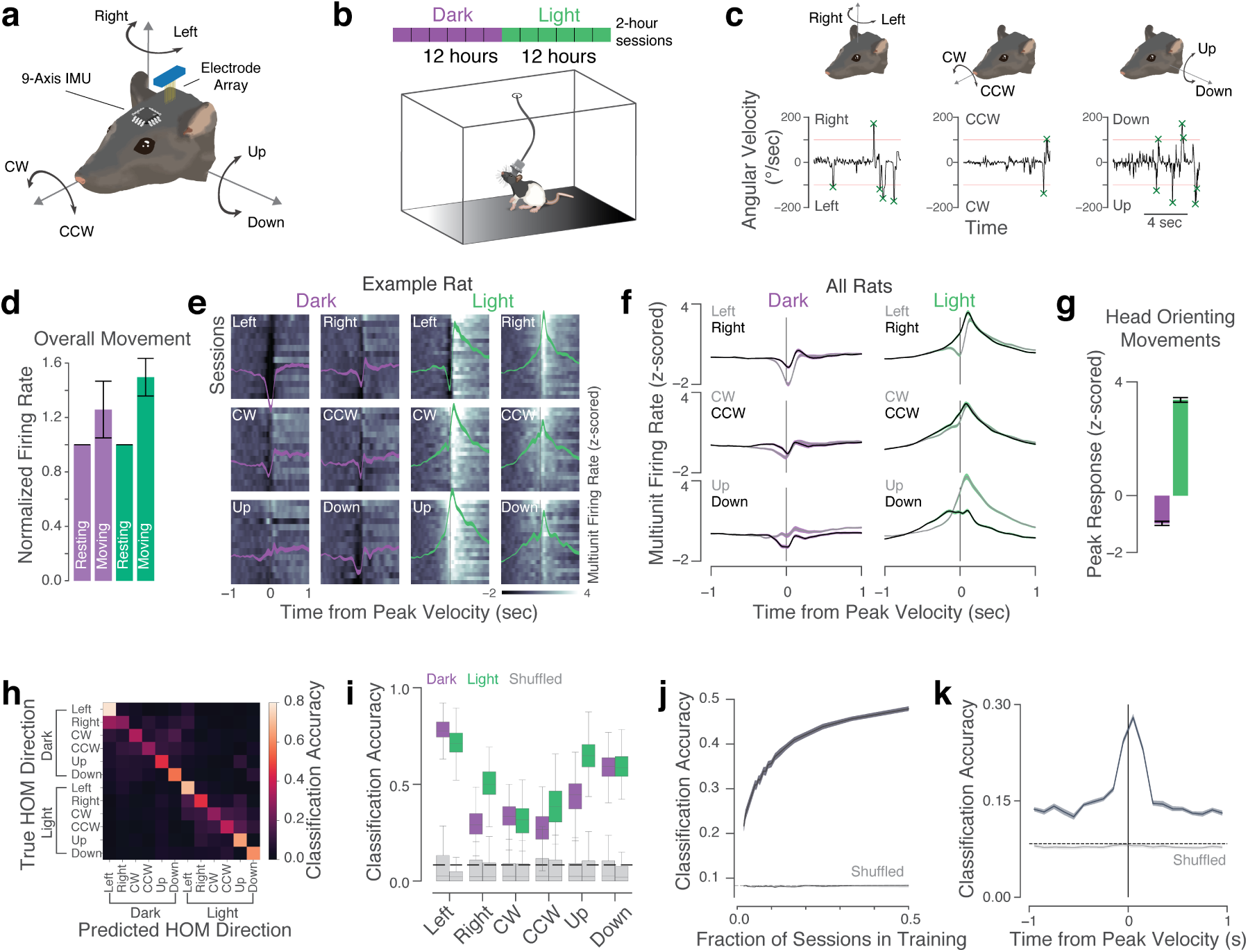
V1 Bidirectional Responses in the Light and Dark Encode Direction of 3D Head Orienting Movements. **(a)** Angular velocity of the head was measured using a head-mounted sensor while neural activity in V1 was measured using chronically-implanted electrode arrays. **(b)** During the 24/7 recordings, rats were free to explore in their home cage either in the light or in the dark, with 12 hours per lighting epoch. Each epoch was then split into 2-hour sessions for analysis. **(c)** Head orienting movements (HOMs) were extracted by finding peaks of the angular velocity signals that crossed a threshold (100°/s). **(d)** Multiunit activity (MUA) firing rates were on average 25.9% and 49.6% higher during overall movement compared to rest, in the dark and light, respectively (*p* < 1*e* − 10 for both, Wilcoxon test; error bars: median absolute deviation). **(e)** Example of z-scored MUA firing rates from one rat’s trial- and tetrode-averaged sessions aligned to extracted left and right (top row), clockwise and counterclockwise (middle row), and up and down (bottom row) HOMs in the dark (purple, first two columns) and light (green, last two columns). Shading in overlaid mean traces represents s.e.m. **(f)** HOM responses averaged across rats, overlaying responses to opposing directions (mean ± s.e.m.). **(g)** In contrast to increased mean firing rates aligned to general movement in **d**, MUA firing rates were suppressed in the dark and enhanced in the light when aligned to HOMs. **(h)** Confusion matrix from logistic regression model trained to predict HOM direction from V1 MUA firing rates. **(i)** Classification accuracy for different HOM directions in the dark (purple), light (green), and for models trained on shuffled data (grey). Dotted line: chance performance (1/12). **(j)** HOM direction decoding performance as a function of the number of sessions used in model training. **(k)** Decoding performance as a function of the lag of the training/testing window relative to peak velocity of the head (100-ms non-overlapping bins).

As previously observed in V1 of head-fixed mice running on spherical treadmills, multiunit activity (MUA) firing rates were indeed higher during overall movement compared to rest, both in the dark and the light (Fig. 1d). In contrast, HOMs were correlated with a sub-second suppression of firing rates in the dark, and an increase of firing rates in the light (Fig. 1e-g). These patterns were consistent across animals (Fig. S2). In the light, V1 activity peaked after peak head velocity, at times roughly corresponding to responses to visual stimulation in rat V1 previously reported (Fig. S1g)^57^. Across all HOMs, V1 activity deviated from baseline as early as 920 ms before movement onset (Fig. S1h), suggesting that these signals reflect a motor or predictive signal rather than sensory reafference. Thus, 3D movement in freely moving animals shapes V1 dynamics.

To address whether observed V1 dynamics encode specific movements, we built a logistic regression classifier that took as inputs the session- and tetrode-averaged MUA in the 500 ms around peak head velocity, and predicted the HOM direction (Fig. 1h,i). The models performed well above chance, with decoding performance higher in the light than in the dark. Further, decoding performance increased as a function of the number of sessions used in training (Fig. 1j). While performance was above chance up to one second before peak velocity, it peaked in the 100 ms after peak velocity (Fig. 1k). Finally, the differences in MUA responses to opposing HOM directions could not be explained by head velocity alone, because the median above-threshold head velocities in opposing directions were indistinguishable from one another (Fig. S1i). These results support the idea that movement direction is encoded in V1, even in the absence of visual input, and that this information is present even before the movements are executed.

### V1 Single Units are Tuned Exclusively to Direction of Movement or Visual Flow

V1 population activity was suppressed in the dark during HOMs in a direction-specific manner. To define the cellular basis for this modulation, we spike-sorted well-isolated units. Sorted single units were classified into regular-spiking putative excitatory units (RSUs) and fast-spiking putative inhibitory units (FSUs) based on waveform shape (Fig. S3). Firing rates of single units were on average increased during epochs of overall movement compared to rest in the dark and the light (Fig. S4a,b). Approximately a fifth of individual RSUs had significantly modulated firing rates during movement, with the vast majority of the significantly-modulated firing rates increasing during movement. A smaller fraction of FSUs had significantly modulated firing rates during movement, although these were more bidirectionally modulated than RSUs (Fig. S4c). RSU mean firing rates were more strongly modulated by the animals’ overall movement than the lighting condition (Fig. S4b), further highlighting the significance of movement on neural coding in V1.

In contrast to the small fraction of units whose mean firing rates changed during overall movement, approximately 75% of single units were responsive to overall movement onset and offset (Fig. S4d). An even larger fraction were responsive during HOMs (Fig. 2a,b). The responses of V1 single units during HOMs were heterogeneous in directionality and timing, with the majority of significantly-modulated units suppressed in the dark and excited in the light (Fig. 2c). While most cells had maximal responses following peak velocity, many peaked before (Fig. 2d). Individual units were specifically tuned to one HOM direction or the other at a rate above chance (Fig. 2e-f), with about half of cells tuned to any particular direction (Fig. 2g).

**Figure 2:**
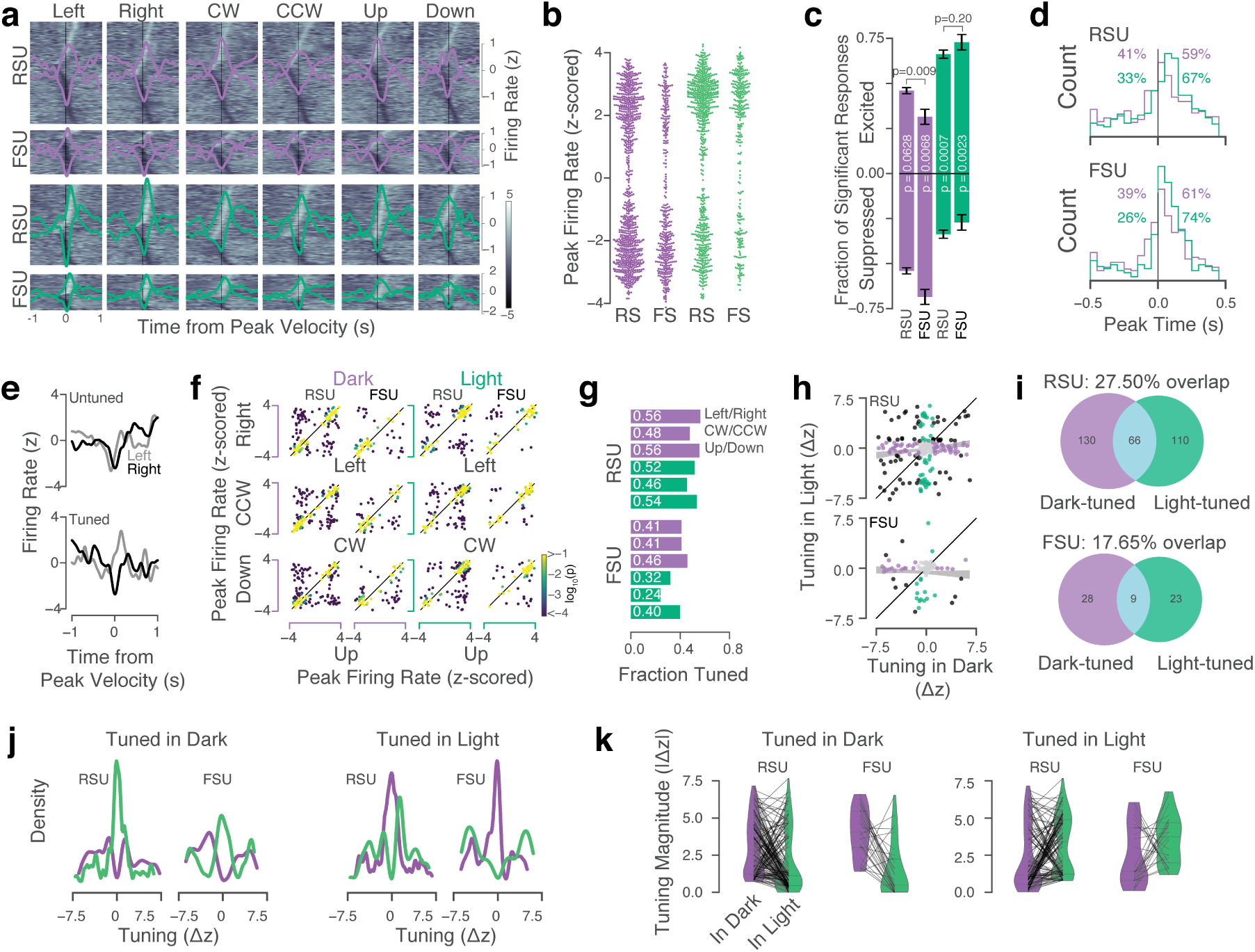
V1 Neurons are Tuned Exclusively to Direction of Movement or Visual Flow. **(a)** Heatmaps of HOM-aligned responses in regular-spiking putative excitatory units (RSUs, *n* = 142 in dark; *n* = 112 in light) and fast-spiking putative inhibitory units (FSUs, *n* = 61 in dark; *n* = 50 in light). Rows represent individual neurons; lines depict means of enhanced and suppressed responses. **(b)** Peak responses of single units to HOMs were bimodal (responses pooled across HOM directions). **(c)** Of the significantly-responding cells, the majority were suppressed in the dark (53.95 ± 1.70% of RSUs and 68.72 ± 4.20% of FSUs) and excited in the light (66.17 ± 2.05% of RSUs and 72.11 ± 3.79% of FSUs) (mean ± s.e.m. across HOM directions; p-values above bars indicate results of t-tests). **(d)** Timing of peak firing rates relative to peak velocity of the head. Responses before peak velocity were more common in the dark than in the light, for both RSUs (*p* = 0.0009) and FSUs (*p* = 0.0006, Fisher’s exact test). **(e)** Two example neurons showing mean responses aligned to left (grey) and right (black) HOMs in the dark. One had similar responses to left and right HOMs (top), while the other was suppressed during right HOMs and excited during left HOMs (bottom). **(f)** Peak responses of neurons (dots) to opposing HOM directions. Similar responses to opposing directions fall along the diagonal. Off-diagonal responses indicate a cell is tuned to HOM direction. Color indicates probability of the responses to the two directions being different from one another (p-value of shuffle test). **(g)** Fractions of significantly-tuned neurons, split by directions (significance at *p* < 0.05, Bonferroni-corrected). **(h)** Direction tuning in dark versus light in units recorded in both lighting conditions (*n* = 100). Green: neurons significantly direction-tuned in the light; purple: neurons significantly tuned in the dark; black: neurons tuned in both conditions; grey: neurons tuned in neither condition. Shading: bootstrap mean ±s.e.m. correlation (RSUs: *r* = 0.13, *p* = 0.048; FSUs: *r* = −0.06, *p* = 0.59). **(i)** Overlap in direction-tuned neurons in dark and light pooling responses across Left/Right, CW/CCW, and Up/Down (*n* = 300 total). **(j)** Left: histograms of direction tuning for neurons tuned in the dark (purple), and the direction tuning of the same neurons in the light (green). Right: same but for neurons direction-tuned in the light (green) and the same neurons’ tuning in the dark (purple). **(k)** Left: absolute tuning magnitude decreases in the light (green) for neurons tuned in the dark (purple). RSU: 3.08 ± 0.15 in dark, 2.04 ± 0.17 in light, *p* = 8*e* − 6; FSU: 4.04 ± 0.30 in dark, 1.44 ± 0.33 in light, *p* = 1*e* − 4, Wilcoxon rank-sums tests. Right: same but for neurons tuned in the light. RSU: 2.23 ± 0.18 in dark, 3.20 ± 0.17 in light, *p* = 1*e* − 4; FSU: 2.43 ± 0.41 in dark, 3.55 ± 0.34 in light, *p* = 0.09, Wilcoxon rank-sums tests.

Consistent with these tuning properties, a logistic decoder trained to discriminate opposing HOM directions (e.g. left vs. right) using single-unit responses performed well above chance (Fig. S5a), with performance exceeding chance prior to movement onset (Fig. S5b), consistent with the possibility that a significant portion of the responses were orienting-based rather than sensory. RSUs and FSUs encoded HOM direction in the dark and light (Fig. S5c-e), with best performance in the light, and mixtures of RSUs and FSUs outperforming models that used only one neuron type. Surprisingly, while RSUs encoded HOM direction better than FSUs in the light, their direction encoding in the dark was slightly poorer than FSUs’ in the dark (Fig. S5d-e). Thus, V1 single unit activity reflected not only overall movement signals (Fig. S4), but HOM direction as well (Fig. 2a-d).

To disambiguate whether direction tuning in the light corresponds to encoding of movement direction, or whether the tuning actually reflects the direction of visual flow, for a subset of our recordings single units were tracked across dark and light sessions. Surprisingly, there was no significant correlation between a cell’s tuning in the dark and in the light (Fig. 2h). Of the cells tuned to HOM direction in either dark or light, fewer than 25% were tuned in both conditions (Fig. 2i). HOM direction-tuned neurons in V1, therefore, are constituted from largely non-overlapping populations.

We reasoned that if V1 contained a neuronal population that discounted the effects of self-generated stimuli on visual processing, that population would exhibit strong strong direction-tuning in the dark and weak tuning in the light. Conversely, if a population encoded the direction of visual flow, it would exhibit strong tuning in the light, but not in the dark. Consistent with this logic, the magnitude of tuning (absolute difference in responses to opposing HOM directions) of cells tuned to HOM direction in the dark was significantly lower in the light, and vice versa, both on average and for most individual units (Fig. 2j,k). Thus, the dynamics of neurons that encoded direction in the absence of visual information in the dark no longer reflected direction when those neurons had access to visual information in the light. Furthermore, neurons that encoded direction while having access to visual- and movement-related information in the light no longer encoded direction when visual information was not available in the dark. Direction-tuned V1 neurons therefore represent two populations that play different functional roles in visual processing: when the lights are on, one population accentuates visual flow direction and another discounts it.

### V1 Responses to Visual Stimulation are Suppressed During Orienting Movements

In freely behaving animals, visual stimuli are processed during both stillness and movement, but the responses of V1 neurons have not previously been examined in unrestrained animals. We wanted to know how responses to visual stimulation would be affected by HOMs, reasoning that if HOMs suppress V1 activity on average, responses to visual stimuli during those movements may be suppressed relative to responses during rest. Alternatively, visual stimulation during HOMs could enhance visual responses, as running does in head-fixed mice^45,47^.

To explore the possibility that HOMs might be associated with altered visual responses, we flashed the behavioral arena lights (on for 500 ms and off for a uniformly random time 400-600 ms) while the rats were behaving freely, and examined average responses during HOMs or rest (Fig. 3a). Flash trials were analyzed according to the animals’ movement in the 100-ms period after stimulus onset, splitting the data into resting or orienting trials (see Materials and Methods).

**Figure 3:**
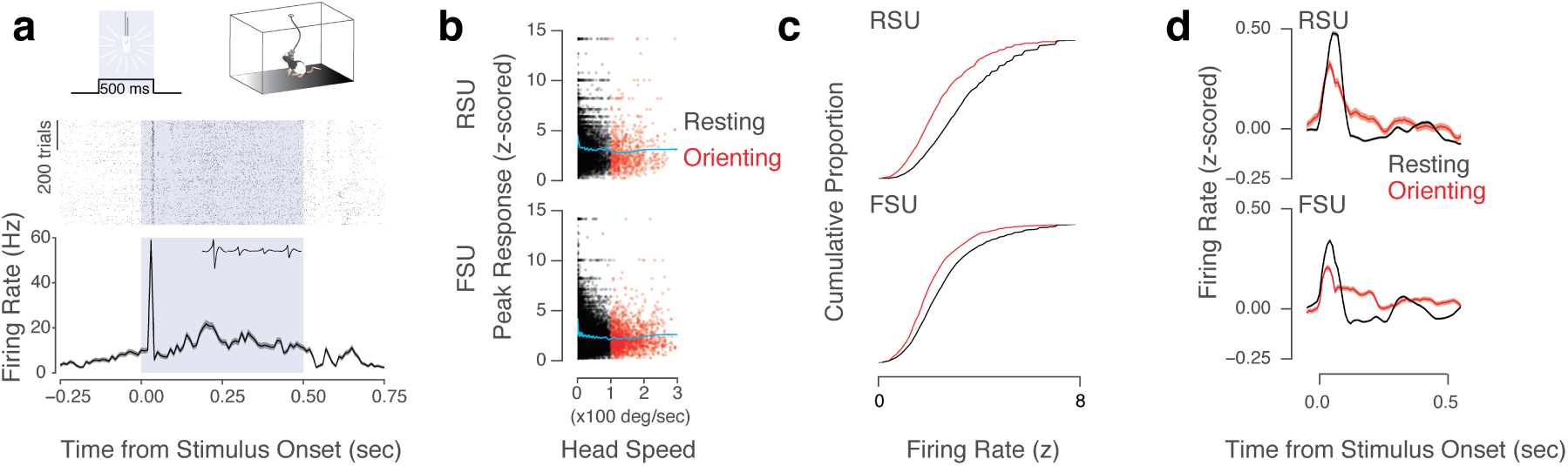
V1 is Suppressed by Visual Stimulation During Head Orienting Movements. Visual stimuli were presented as 500-ms flashes of the overhead cage lights over the course of several hundred trials. **(a)** Response profile of one V1 cell over *n* = 707 trials, with spike raster plot (middle) and mean ± s.e.m. response (bottom). Purple shading represents stimulus timing. Inset: waveform of the example unit. **(b)** Peak responses to flash stimuli as a function of head speed. Trials are colored by movement condition: resting (black) or orienting (red), with orienting defined as the angular velocity of the head exceeding 100°/sec. Each dot is one trial (*n* = 9692 trials from *n* = 52 RSUs and *n* = 20439 trials from *n* = 52 FSUs from *n* = 12 sessions with *n* = 656.58 ± 135.76 trials each, across *n* = 3 rats). Cyan lines indicate the rolling mean. **(c)** Cumulative proportion of peak firing rates in the response window. **(d)** Mean responses of RSUs (top) and FSUs (bottom) in resting or orienting conditions. Responses were lower during HOMs in RSUs (*p* = 4.4*e* − 19 MWU test) and FSUs (*p* = 1.8*e* − 30 MWU test)

Responses to flashes were largest when the animals were resting and were suppressed during HOMs, to a greater extent in excitatory than inhibitory cells (Fig.3b-d). While peak responses to visual stimulation during HOMs were lower than during rest, the activity also persisted longer (Fig.3d). The finding that V1 visual responses are suppressed during orienting movements supports the idea that the rodent visual system has a mechanism to cancel the effects of movements on sensory perception, akin to previous findings in rodent auditory cortex^27^ and primate visual areas^10–14^.

### V1 Head Orienting Responses Depend on Secondary Motor Cortex

The observation that V1 responded to HOMs in the dark before HOM onset suggests that HOM-related signals in V1 might originate in a motor region of the brain. A multitude of anatomical, functional, and electrophysiological evidence pointed to secondary motor cortex as a likely source of HOM signals in V1. We therefore lesioned large portions of M2 (including the area that projects to V1) via bilateral injections of ibotenic acid (Fig. 4a-c; S6; Fig. S7a). The lesions did not alter orienting behaviors (Fig. 4d; Fig. S7b,c); further, MUA firing rates were still higher during overall movement compared to rest, albeit to a lesser extent compared to non-lesioned animals (Fig. S7d). V1 activity in lesioned animals responded to visual stimulation (Fig. S7e-k), but exhibited significantly reduced HOM-related responses (Fig. 4e). These trends were consistent across animals and HOM directions (Fig. S8a). Across all HOM directions, the previously observed negative responses in the dark and positive responses in the light were largely diminished in lesioned animals, with the magnitude of responses in the dark reduced more so than the magnitudes of responses in the light (Fig. 4f). HOM direction encoding was greatly reduced in models trained and tested on lesioned animals’ data (Fig. 4g), with most dramatic drops in performance for Left, Right, CW, and CCW directions in lesioned animals both in dark and light (Fig. S8b). M2 therefore plays a crucial role in shaping HOM-related responses and HOM direction information in V1.

**Figure 4:**
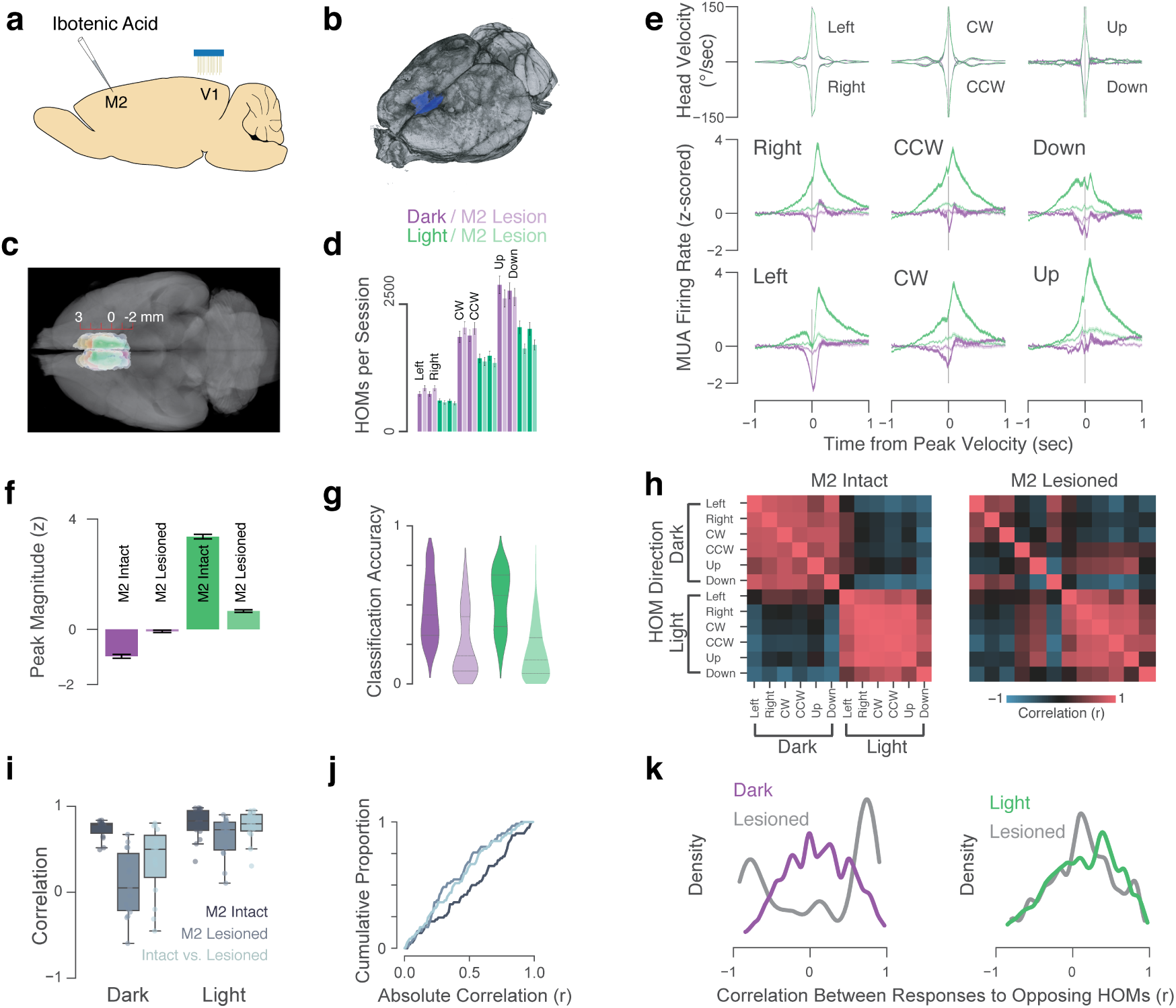
Head Orienting Movement Signals in V1 Depend on Secondary Motor Cortex. **(a)** M2 was lesioned bilaterally using injections of excitotoxic ibotenic acid. Tetrode arrays were implanted in V1 following lesioning. **(b)** An example lesioned brain imaged using micro-CT. Lesion area highlighted in blue. **(c)** Horizontal projection of lesion ROIs (*n* = 4 rats) overlaid over an example brain. Anterior-posterior coordinates relative to Bregma. **(d)** Numbers of HOMs extracted per session from non-lesioned (darker purple and green) and lesioned rats (lighter purple and green) were indistinguishable from one another (all *p* > 0.02, Mann-Whitney U tests on lesioned versus non-lesioned numbers). **(e)** Mean HOM-aligned angular velocities of the head and MUA firing rates across animals and conditions. Top row: extracted HOMs (velocity). Middle and bottom rows: z-scored MUA firing rates averaged across *n* = 5 non-lesioned animals, *n* = 4 lesioned animals, *n* = 16 tetrodes each. **(f)** Peak MUA magnitudes in the 0.5 seconds around peak velocity were significantly lower in lesioned rats in the dark (*p* < 1.5*e* − 41, MWU test) and light (*p* < 3.9*e* − 109, MWU test). **(g)** HOM direction decoding models trained and tested on lesioned animals performed poorly (classification accuracy: 0.26 ± 0.01 for dark; 0.19 ± 0.01 for light) compared to non-lesioned models (0.45 ± 0.01 for dark; 0.53 ± 0.01 for light) (mean ± s.e.m.). **(h)** Correlation structure of MUA traces among HOM directions in dark and light in non-lesioned (left) and lesioned (right) rats. **(i)** Correlations are reduced in lesioned animals, more so in the dark (*r* = 0.73 ± 0.03 in non-lesioned and *r* = 0.10 ± 0.10 in lesioned rats *p* = 0.00065, MWU test) than in the light (*r* = 0.80 ± 0.05 in non-lesioned and *r* = 0.65 ± 0.06 in lesioned rats *p* = 0.00065, MWU test). **(j)** Cumulative proportions of absolute correlations among HOM directions. **(k)** Correlation of single-unit responses during opposing (Left vs Right, CW vs CCW, Up vs Down) HOM directions. In the dark, 49.0% of units in lesioned rats highly correlated responses (*r* > 0.5), compared to 20% in non-lesioned rats; and 31.4% had highly anti-correlated responses (*r* < −0.5) compared to 6.2% in non-lesioned rats. Conversely, in the light, 18.7% of units in lesioned rats highly correlated responses (*r* > 0.5), compared to 23.5% in non-lesioned animals; 8.1% of units in lesioned rats had highly anti-correlated responses, similar to the 7.4% of units in non-lesioned animals.

The reduced HOM direction information in lesioned animals led us to compare the correlation structure among the average MUA traces in lesioned and non-lesioned animals (Fig. 4h-j; see Materials and Methods). MUA responses to HOMs in lesioned rats were reduced in the dark, but remained high in the light (Fig. 4i,j). Single unit correlations in non-lesioned animals were broadly distributed both in the dark and light, reflecting a capacity to encode HOM direction (Fig. 4k). In contrast, in lesioned animals, over half of the neurons had highly correlated responses to opposing HOM directions in the dark (compared to 20% in non-lesioned animals), and one third were highly anticorrelated (compared to 6% in non-lesioned animals). Correlation structure in the light appeared to be qualitatively similar between lesioned and non-lesioned animals, likely reflecting common visual drive. These results demonstrate that M2 shapes responses to orienting movements of the head in V1, suggesting that M2 may support a neural mechanism that predicts the direction of visual flow given a motor command and cancels the effects of such flow on sensory processing.

## Discussion

Vision has largely been studied in a passive context, with restrained animals passively absorbing stimuli projected on a screen. In their natural states, however, animals interact with sensory stimuli of various modalities by sniffing, whisking or palpating, or making saccadic movements of the eyes, head, or body. Saccadic eye movements in primates have been studied extensively, but while rodents also make complex patterns of movements of the head and eyes, their functional purpose and impact on visual processing is for the most part unknown. We reasoned that recording neuronal activity in V1 of freely behaving rats could help clarify the impact of naturalistic movements on sensory cortical dynamics. We found that V1 dynamics in rats during free movement precisely relate to 3D movements of the head in a manner that depends on secondary motor cortex. V1 multiunit firing rates were suppressed during orienting movements of the head in the dark, and enhanced in the light on a sub-second basis. Further, around half of individual neurons were tuned to particular HOM directions in a manner that depended on the presence or absence of light. Our data therefore suggest a mechanism through which cortex can distinguish self-generated sensory signals and those that come from the outside world; these findings further raise the possibility that orienting movements of the head serve the same functional purpose in rodents as saccadic eye movements do in primates.

Movement-related neuronal activity has been found across virtually all regions of the mammalian brain, but its function remains unknown^55,56^. Locomotion signals in V1 have been proposed to predict visual flow direction^40,54^; such signals could support a predictive coding framework in which V1 computes deviations from expected flow, thereby making long-range communication among visual areas more efficient^49,58,59^. A related possibility is that movement signals in sensory cortex could reflect efference copy signals from motor regions. Such movement signals have been identified in the rodent auditory system and other sensory systems^13,14,27,60–63^. While a signal predicting visual flow and an efference copy of movement plans are similar in nature, the latter could be used to cancel the effects of purposive movement on sensory processing by suppressing activity during movement. Our observation that V1 multiunit and the majority of single units were suppressed in the dark suggests that V1 dynamics reflect not just a prediction of visual flow, but also a cancellation of the effects of flow on sensory processing. Further, the fact that V1 dynamics were modulated in an HOM direction-specific manner suggests that the cancellation reflects specific 3D movement dynamics.

Our experiments identify two functional populations: one that is direction-tuned in the dark and cancels out the effects of movement-related signals on sensory responses, and one that is direction-tuned in the light and encodes direction of visual flow. Further, responses of V1 single units to visual stimulation were diminished during HOMs compared to rest. These data are reminiscent of the suppression of activity observed in auditory cortex during locomotion, but stand in contrast with the enhanced neural responses observed in visual cortex during locomotion^45,47,48,64,65^. One possible explanation of this discrepancy is that running and orienting movements produce opposing effects on visual responses, perhaps because of differing consequences on visual flow (i.e. translation vs rotation).

Lesions to M2 severely diminished HOM-related activity while sparing the orienting behaviors (Fig. 4) and preserving responses to visual stimulation (Fig. S7d-i). While M2 axons project directly to V1 (Fig. S6)^40^, it is possible that HOM-related activity in V1 is not directly inherited from axonal projections from M2, but from M2’s influence on retrosplenial cortex (RSC)^54^ or thalamic nuclei^50^. It is also possible that V1 inherits HOM-related signals from the LGN, which has been shown to be affected by locomotion^52^. Further elucidation of the movement signal pathways to V1 will require genetic tools to silence or record the dynamics of M2-projecting axons over V1 in a freely behaving context^40^.

A major challenge for studying vision in unrestrained animals is the control of visual stimulation, a problem that might be solved using head-mounted LED screens or cameras^17^. Recent work in head-fixed mice running in virtual reality (VR), where visual flow is yoked to locomotion, represents an important advance in studying vision as an active sensation^40,49,66,67^. Still, most VR setups require head restraint, which prevents naturalistic interaction with visual stimuli.

Our data show that various orienting movements in 3D have profound effects on neuronal dynamics in V1, a cortical area traditionally thought to primarily perform feed-forward computations on incoming visual information. While the 3D movements examined here are closer to naturalistic behavior than locomotion in restrained animals, the characterization of free movement as HOMs along three axes is still lacking compared to the complex patterns of movement animals actually perform; as such, it will be critical to continue examining sensory cortical dynamics using computational neuroethology methods that reveal the structure of naturalistic behavior^68^.

Saccadic eye movements in primates serve to shift gaze, foveate, and explore a visual scene. These movements are accompanied by a suppression of activity in visual brain areas, which serves to reduce blur. Rodents make complex patterns of head and eye movements, some of which are used to estimate distances, but whether or not these also serve to explore a visual scene–a retinal palpation–is not clear because the rodent retina does not have a fovea. Our finding that V1 activity is suppressed during orienting movements of the head, just as primate visual areas are suppressed during saccades, raises the intriguing possibility that such orienting movements serve the same functional purpose in rats that saccades do in primates. Our findings are consistent with a model in which sensation and action are deeply intertwined processes: animals actively sample the sensory world in order to make decisions and learn. Understanding vision therefore will be facilitated by considering it an active process, and exploring its underlying neural codes while subjects are free to behave.

## Materials and Methods

### Animals

The care and experimental manipulation of all animals were reviewed and approved by the Harvard Institutional Animal Care and Use Committee. Experimental subjects were female Long Evans rats 3 months or older, weighing 300-500 g (*n* = 10, Charles River, Strain Code: 006).

### Surgery

Rats were implanted with 16-tetrode electrode arrays targeting L2/3 of V1. Animals were anesthetized with 2% isoflurane and placed into a stereotaxic apparatus (Knopf Instruments). Care was taken to clean the scalp with Povidone-iodine swabsticks (Professional Disposables International, #S41125) and isopropyl alcohol (Dynarex #1204) before removing the scalp and cleaning the skull surface with hydrogen peroxide (Swan) and a mixture of citric acid (10%) and ferric chloride (3%) (Parkell #S393). Three to four skull screws (Fine Science Tools, #19010-00) were screwed into the skull to anchor the implant. A 0.003” stainless steel (A-M Systems, #794700) ground wire was inserted ∼2mm tangential to the brain over the cerebellum.

Tetrodes were arrayed in 8×2 grids with ∼250-micron spacing, and were implanted in V1 with the long axis spread along the AP (ranging 6-8 mm posterior to bregma, 4.5 mm ML, targeting layer 2/3 at 0.6 mm DV). The dura was glued (Loctite) to the edges of the craniotomy to minimize movement of the brain relative to the electrodes. After electrodes were inserted into the brain, the craniotomy was sealed with Puralube vet ointment (Dechra) and the electrodes were glued down with Metabond (Parkell). Post-operative care included twice-daily injections of buprenex (0.05mg/kg Intraperitoneal (IP)) and dexamethasone (0.5 mg/kg IP) for three days.

### Behavior

Spontaneous behavior in rats living in a 15×24” home cage was recorded under three conditions: Dark, in which the lights in the box and room were turned off; Light, in which the box was illuminated; and Flash, in which visual responsiveness of neurons was assessed by flashing the lights repeatedly (On: 500 ms; Off: 400-600 uniformly random time). Recordings were carried out 24/7 and split into ∼12-hour Dark and ∼12-hour Light sessions, with 10-30-minute Flash sessions interspersed in a some of the experiments.

The behavior box was constructed from aluminum extrusions and black extruded acrylic (McMaster). The floor was covered in bedding and the arena contained a cup with food, a water bottle and toys. The walls were lined with strips of white tape at different orientations to provide visual features in the Light condition, and the box was outfitted with white LED strips (Triangle Bulbs Cool White LED Waterproof Flexible Strip Light,T93007-1, Amazon) to provide illumination. For the Dark condition recordings, room and box lights were turned off and care was taken to make sure most sources of light in the experimental room (e.g. LEDs from computers and other hardware) were covered with black tape. To assess whether the recording box was sufficiently dark, a test was performed with a human subject (GG) acclimated in the room for 30 minutes, after which visual features in the cage were still not visible.

For recordings, rats were tethered with a custom 24” cable (Samtec, SFSD-07-30C-H-12.00-DR-NDS, TFM-107-02-L-D-WT;McMaster extension spring 9640K123) to a commutator (Logisaf 22mm 300Rpm 24 Circuits Capsule Slip Ring 2A 240V TestEquipment, Amazon). A 9-axis Inertial Measurement Unit (IMU) (BNO055, Adafruit) was used to record movement; the sensor was epoxied to the connector on the cable, in a way that placed it directly above the electrodes and headstage. This not only ensured that the sensor was always in the same position above the animals’ heads, but also that it stayed powered after the animals were unplugged, preventing the need to re-calibrate the sensor after each recording. The IMU data were acquired at 100Hz using a micro-controller (Arduino) and saved directly to the acquisition computer’s disk. To synchronize IMU and electrophysiology data, the Arduino provided a 2-bit pseudo-random pulse code to the TTL inputs on the electrophysiology system.

The 9-axis IMU combines signals from a 3-axis accelerometer, 3-axis magnetometer, and 3-axis gyroscope. It is equipped with a data-fusion algorithm that combines the signals to calculate absolute direction in three axes, yielding a vector of yaw, roll, and pitch; in addition, the sensor outputs linear acceleration in the forward/backward, left/right, and up/down directions. The derivatives of the yaw, roll, and pitch signals (which reflect left/right CW/CCW, and up/down angular velocities of the head, respectively), were used to assess V1 dynamics during HOMs. To extract HOMs, peaks in crossings of a 100 deg/sec threshold of the angular velocity traces were found. The L2 norm of the linear acceleration components was used as a proxy for the overall movement.

### Electrophysiology

Tetrodes were fabricated using 12.5-micron nichrome wire (Sandvik-Kanthal) following standard procedures^69–71^. Tetrodes were threaded through 42 AWG polyimide guide tubes into 8×2 grids of 34 AWG tubes (Small Parts) and glued to a single-screw micro-drive. The drive was modified from a design in Mendoza et. al^72^ and Vandercasteele et. al^73^, in which a 3-pin 0.1” header served as the skeleton of the drive, with a #0-80 screw replacing the middle pin, and the headers plastic serving as the shuttle. In the experiments reported here, the tetrodes were not advanced after recording sessions started. The tetrodes were plated with a mixture of gold (Neuralynx) and polyethylene glycol (PEG) as per Ferguson et. al^74^, to an impedance of ∼100 − 250*K*Ω. The ground and reference wires were bridged and implanted through a craniotomy above the cerebellum.

Electrode signals were acquired at 30 kHz using custom-made Intan-based 64-channel headstages^75^ and Opal-Kelly FPGAs (XEM6010 with Xilinx Spartan-6 ICs). Spikes were extracted following procedures described in Dhawale et al (2017)^75^. Multiunit firing rates were estimated in non-overlapping 10-ms bins from extracted spikes. Multiunit and single-unit firing rates were Gaussian-filtered and in some cases z-scored. Single-units were sorted using MountainSort^76^ and classified into putative excitatory regular-spiking units (RSUs) or putative inhibitory fast-spiking units (FSUs) based on the trough-to-peak time (width) and full-width at half-max (FWHM) of the unfiltered waveforms (Fig. S3).

### Viral Tracing

In order to localize the V1-projecting portion of M2, viral tracing experiments between these two regions were performed using retrograde viruses and with conditional expression of fluorophores (Fig. S6). retro-AAV-GFP (Janelia) and retro-AAV-Cre (AAV pmSyn1-EBFP-Cre; Addgene #51507) were injected into three sites in V1, at 6.35 mm anterior-posterior axis (AP), 4.55 mm mediolateral (ML), 6.0 AP, 4.47 ML, and 5.8 AP, 4.5 ML, at 1 mm below the brain surface, with 500 nl at each site, injected at 25 nl/min using an UMP3 UltraMicroPump (WPI). Into putative V1-projecting portion of M2, FLEX-tdTomato (AAV-CAG-FLEX-tdTomato, UNC Vector core) was injected at two sites (0.5 mm AP, 1.0 mm ML, 1.8 mm DV; −0.5 AP, 0.9 ML, 1.8 and 0.8 DV; 200 nl each at 25nl/min). With this strategy, we were able to visualize retrogradely-labeled inputs to V1 in green and V1-projecting M2 axons in red, imaged on slices stained for GFP (Rabbit anti-GFP, Life Technologies A11122, 1:1000; Alexa Fluor 488 Goat Anti-Rabbit, Life Technologies A-11034, 1:1000) and for RFP (Chicken Anti-RFP, Sigma, AB3528, 1:500; Alexa Fluor 568 Goat Anti-Chicken IgG, Life Technologies, A-11041, 1:1000) using an Axio Scan.Z1 slide scanner (Zeiss).

### Lesions

Lesions of M2 were performed using excitotoxic injections of ibotenic acid (IA) (Abcam ab120041) delivered using an UMP3 UltraMicroPump (WPI) during two separate procedures. Aliquots of IA were prepared at 1% concentration and frozen. In the first procedure, IA was injected into four sites in one hemisphere (1.5 mm AP, relative to Bregma and 1.0 mm ML; 0.5 AP, 0.75 ML; −0.5 AP, 0.75 ML; and −1.5 AP, 0.75 ML, with two injections per site, at 1.6 and 0.8 mm below the brain surface, 75 nl each) and the animal was allowed to recover for one week, after which the injections were repeated at the same sites in the opposite hemisphere and electrode arrays were implanted in V1.

### Micro-CT

X-ray micro-computed-tomography (micro-CT) was used to quantify lesion sizes and locations and to determine the depth of implanted electrodes after performing electrolytic lesions through the implanted tetrodes by passing 40*µA* through each wire for 15 seconds^77^. To extract brains, rats were deeply anesthetized with sodium pentobarbital (180 mg/kg; Fatal-Plus C IIN, Vortech Pharmaceuticals, Dearborn, MI) and perfused with paraformaldehyde (#15710, Electron Microscopy Sciences (EMS), Hatfield, PA) and glutaraldehyde (#16220, EMS). Brains were then stained with osmium (VWR 19190) for two weeks and embedded in resin before imaging using a Nikon Metrology X-Tek HMX ST 225 Micro-CT scanner (Nikon Metrology Ltd., Tring, UK). This method, which was previously described in detail in Masis et. al^77,78^, allows characterization of lesions without slicing the brain. Lesion ROIs from scanned brains were analyzed using FIJI^79^ and ITK-SNAP^80^.

### Head Orienting Movement Extraction

HOMs were extracted from the time derivatives of head direction signals (yaw, roll, and pitch) by finding crossings of a threshold (100 deg/sec). Consecutive threshold crossings were peak-aligned. Positive and negative crossings of the threshold in the derivative of yaw signal were termed Left and Right HOMs, respectively; similarly, roll crossings were termed clockwise (CW) and counterclockwise (CCW) HOMs, and pitch crossings were termed Up and Down HOMs.

### Movement Classification

In addition to HOMs, rats were classified as moving or resting (Fig. S1d,e) based overall movement, which was calculated as the L2 norm of the linear (translational) components of acceleration. Histograms of the overall movement signal showed a bimodal distribution (Fig. S1d,e). The valley between the two peaks was used as the threshold at which the animals were considered to be immobile (resting) or mobile (moving)^75^.

### Head Orienting Movement Direction Decoding

HOM direction was decoded using MUA firing rates (Fig. 1) or single-unit firing rates (Fig. 2; Fig. S5) using multinomial logistic regression implemented in the Scikit Learn Python package^81^. In the MUA decoding, model inputs consisted of z-scored firing rates averaged across tetrodes (*n* = 16) and sessions (*n* = 86 − 107) in a 0.5-sec window centered at peak velocity (Fig. 1h-j) or a 0.1-sec sliding window −1 to +1 sec around peak velocity (Fig. 1k). The models were trained and tested on half (Fig. 1h,i) or varying fractions (Fig. 1j) of the sessions. SUA decoding models were constructed similarly, but with each neuron’s firing rate contributing a separate feature to the model. The models were trained either to decode opposing HOM directions (e.g. Left vs Right; CW vs CCW; Up vs Down) (Fig. S5a,b), or six-direction classification (Fig. S5c-e). The latter were trained either on varying numbers of HOM bouts (up to *n* = 100 trials) using all neurons (Fig. S5c); or with half (*n* = 50) of the trials used for training and the other half for testing, using varying numbers of neurons (Fig. S5d,e). All models were trained with *n* = 100 random splits of trials or neurons. Each model was threefold cross-validated to find the optimal regularization parameter C (ranging from 1*e* − 03 to 1*e* + 03 and penalty (L1 or L2).

### Flash Stimulus Presentation and Analysis

Visual stimuli consisted of 500-ms flashes of white LEDs mounted on the ceiling of the home-cage arena, which was fully dark during the flash off-cycle (400-600 ms, uniform random time).

Responses to flash stimuli were quantified in the 100-ms window following stimulus onset and split into two conditions according on the rat’s movement parameters within that window: orienting, in which the angular velocity of the head crossed the 100 deg/sec threshold in either of the three axes; or resting, in which the angular velocity and total acceleration were below their thresholds. Post-stimulus time histograms (PSTHs) were smoothed with a Gaussian filter and z-scored.

Responsive cells were manually selected by examining waveforms and mean PSTHs, following which significant responses were selected by discarding trials with zero variance and performing a shuffle test between mean firing rates in the response window (50-ms between 30 and 80 ms after stimulus onset) and a baseline window (50-ms before stimulus onset). Responses during movement and rest were compared using Mann-Whitney U test in the response window (Fig 3d).

### Statistics

All statistical comparisons were done using non-parametric tests (e.g. Mann-Whitney U or Wilcoxon) unless specified otherwise. A significance level of alpha = 0.01 was used throughout, unless otherwise noted. Bonferroni correction was applied where appropriate.

## Acknowledgements

We would like to thank Jeffrey Markowitz, Bence Ölveczky, Sandeep Robert Datta, and Juliana Rhee for advice and helpful comments on the manuscript. We thank Gerald Pho for providing ibotenic acid and advice on lesion segmentation. Edward Soucy, Brett Graham, and Joel Greenwood of the CBS Neuroengineering Core were instrumentally helpful in technical advice and on light measurements. Douglas Richardson of the Harvard Center for Biological Imaging helped with slice imaging. H. Greg Lin of the Harvard Center for Nanoscale Systems assisted with micro-CT imaging. This work was performed in part at the Center for Nanoscale Systems (CNS), a member of the National Nanotechnology Coordinated Infrastructure Network (NNCI), which is supported by the National Science Foundation under NSF award no. 1541959. CNS is part of Harvard University. GG was supported by the National Science Foundation (NSF) Graduate Research Fellowship Program (GRFP).

## Author contributions statement

GG conceived and performed the electrophysiology experiments, analyzed the data, and wrote the manuscript. SBEW designed the viral tracing and lesion experiments. JM designed and performed the micro-CT and osmium staining experiments. DC provided research funding and space.

## Additional information

### Competing Interests

None.

## Supplementary Figures

**Figure S1:**
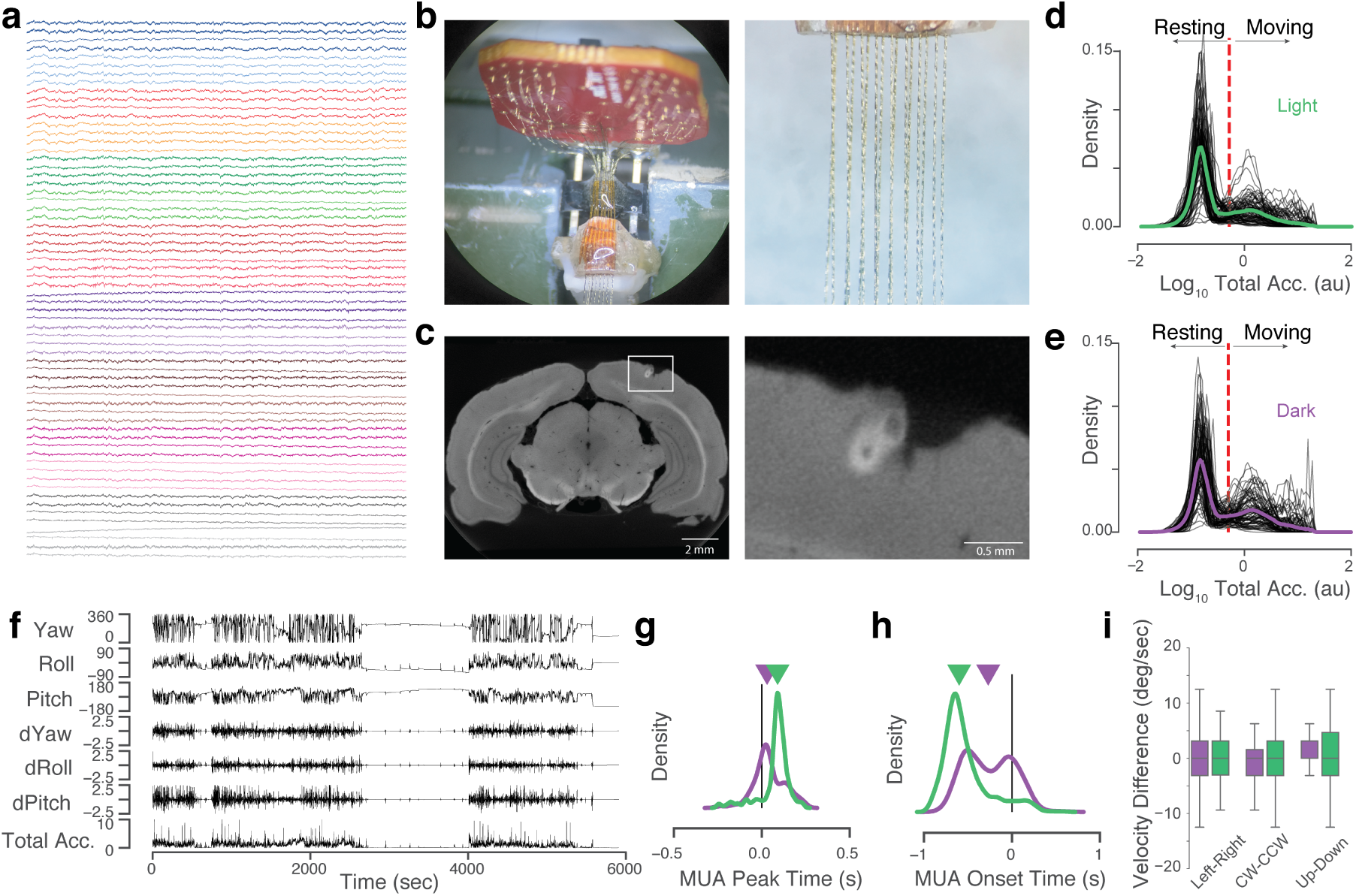
(Related to Figure 1). Chronic Recording of Neural Activity in L2*/*3 of V1 and Head Velocity Signals. **(a)** Unfiltered traces (1 sec) from the 64 channels of the 16-tetrode array colored by tetrode. **(b)** 16-tetrode array design (left) and closeup of the tetrodes (right). **(c)** Micro-CT digital coronal slice (left) showing electrolytic lesions over the implant site in V1 (∼-6.00 mm AP) marking tetrode locations. Mean implant depth: ∼350 microns below the surface (L2/3). Right: detail of white box in left panel. **(d-e)** Classification of behavior into ‘resting’ or ‘overall movement’ based on total acceleration (the L2 norm of the linear components of acceleration) in light (**d**) or dark (**e**). Probability density plots of total acceleration (black: individual sessions; green/purple: mean across sessions) show a bimodal distribution. Dotted red lines show manually-selected classification boundary. **(f)** Example behavioral recording showing yaw, roll, and pitch angles (degrees), their derivatives (x100 deg/sec), and the total acceleration. **(g)** Time of peak MUA firing rates relative to peak angular velocity of the head in the dark (0.0423±0.0038) and light (0.0741±0.0036) (*p* = 2.8*e*−19, MWU test). **(h)** Time of MUA deviation from baseline relative to peak angular velocity of the head (dark: −0.2552±0.0117 (s) (mean± s.e.m.), light: −0.5261 ± 0.01109 (s), *p* = 2.8*e*−61, MWU test). Carets indicate medians. **(i)** Differences in median above-threshold velocities across sessions and rats for opposing HOM directions are centered around zero, indicating that asymmetries in velocities cannot explain differential MUA responses to various HOM directions (Wilcoxon test of Left-Right velocity differences *p* = 0.59 in dark and *p* = 0.67 in light; CW-CCW *p* = 0.51 in dark, *p* = 0.92 in light; and Up-Down *p* = 0.0042 in dark, and *p* = 0.13 in light).

**Figure S2:**
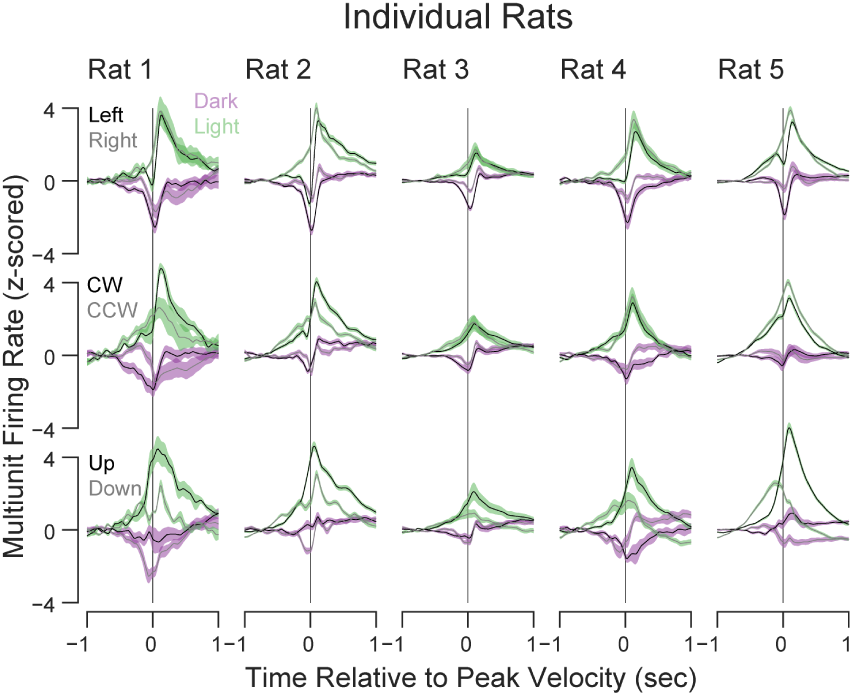
(Related to Figure 1). Head Orienting Movement Responses in Individual Rats. Mean (± s.e.m.) MUA firing rates (z-scored) during Left/Right (top row), CW/CCW (middle row), and Up/Down (bottom row) HOMs in the dark (purple) and light (green) averaged across tetrodes and HOM bouts. As in Fig1e-f but showing individual animals (columns).

**Figure S3:**
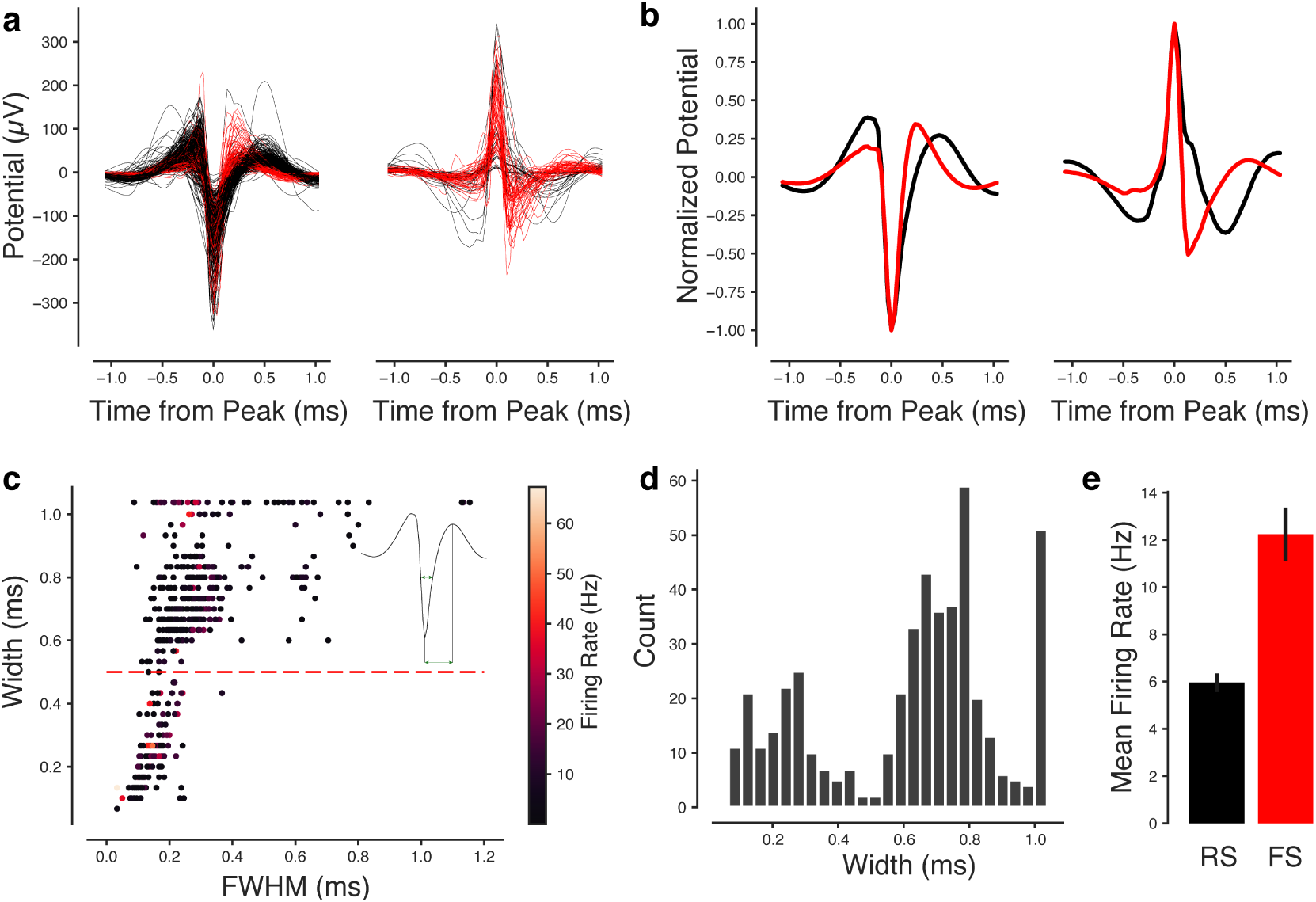
(Related to Figure 2). Single-Unit Waveform Classification. **(a)** Mean waveforms of *n* = 365 single units recorded in dark (*n* = 203) or light (*n* = 162) in *n* = 5 rats across *n* = 10 sessions, separated by negative-peaked (left) and positive-peaked (right) units. Waveforms were taken from the highest-amplitude channels within a tetrode. Color indicates fast-spiking units (FSUs, red) and regular-spiking units (RSUs, black) based on classification in **c. (b)** Peak-normalized waveforms averaged across units. **(c)** Spike width (trough to peak time) vs. full-width-at-half-max (FWHM) calculated for each unit’s unfiltered waveform. Each dot is a unit, colored by its mean firing rate. Inset: mean waveform illustrating width and FWHM calculation. Dotted line: classification boundary for fast-spiking putative interneurons (FSUs; width<0.5ms) and regular-spiking putative excitatory cells (RSUs; width>0.5ms). **(d)** Histogram of mean spike widths. **(e)** Mean firing rates of RSUs and FSUs. Red: FSUs (*n* = 61 in Dark; *n* = 50 in Light), Black: RSUs (*n* = 142 in Dark; *n* = 112 in Light).

**Figure S4:**
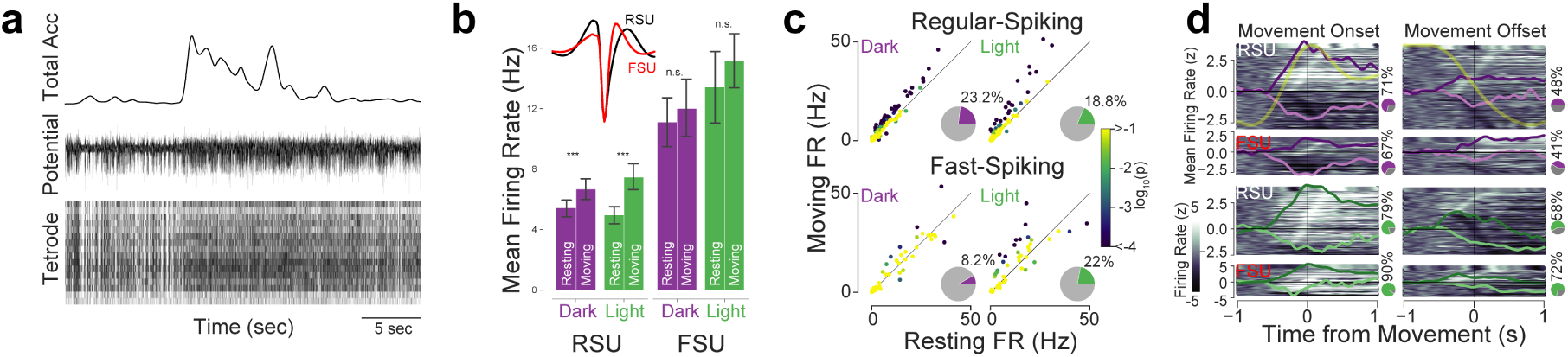
(Related to Figure 2). Single-Unit Activity During Overall Movement. **(a)** Example bout of movement (top), quantified as overall movement (total acceleration exceeding movement threshold set in Fig. S1) and aligned to a raw bandpass-filtered trace from one electrode (middle) and extracted multi-unit spikes (bottom). **(b)** Mean firing rates of single-units during overall movement or rest, in light (green) or dark (purple), separated by unit type (regular-spiking (RSU) putative pyramidal cells and fast-spiking (FSU) putative interneurons). Inset: mean waveforms of RSUs and FSUs. Mean RSU firing rates were higher during overall movement compared to rest (dark, *p* = 1.45*e* − 7; light, *p* = 5.13*e* − 16, Wilcoxon rank-sums tests), whereas mean FSU firing rates were higher but not statistically significantly affected by overall movement (dark, *p* = 0.19; light, *p* = 0.046). Mean firing rates were not affected by lighting condition in RSUs during overall movement (dark vs light, *p* = 0.22) or rest (dark vs light, *p* = 0.16), or in FSUs during movement (dark vs light, *p* = 0.02) or rest (dark vs light, *p* = 0.06). Mean firing rates were thus more strongly modulated by overall movement than lighting condition. **(c)** Firing rates of individual units during overall movement and rest. RSU firing rates were higher during overall movement in dark and light, whereas firing rates of individual FSUs were either higher or lower. Inset pie charts: percentages of units with firing rates modulated by overall movement. Out of 142 RSUs recorded in the dark, 33 were significantly modulated by overall movement (23.2%; permutation test; *p* < 0.05, Bonferroni-corrected), while 5 out of 61 FSUs were modulated in the dark (8.2%). In the light, 21 of 112 (18.8%) RSUs and 11 of 50 (22%) FSUs were modulated by overall movement. Each dot represents one neuron; color represents p-value in permutation test. **(d)** Firing rates aligned to overall movement onsets and offsets. Heatmaps showing z-scored firing rates of individual units (y-axis) over a 2-second period (x-axis) aligned to overall movement onsets (left column) or offsets (right). Lines show the mean overall movement (yellow) and mean z-scored firing rates (purple: dark; green: light; means are separated by sign of activity in the 0.5 seconds around movement detection (darker traces: positive responses; lighter: negative). Pie charts show percentages of units significantly modulated by overall movement onsets or offsets (permutation test; *p* < 0.05, Bonferroni-corrected).

**Figure S5:**
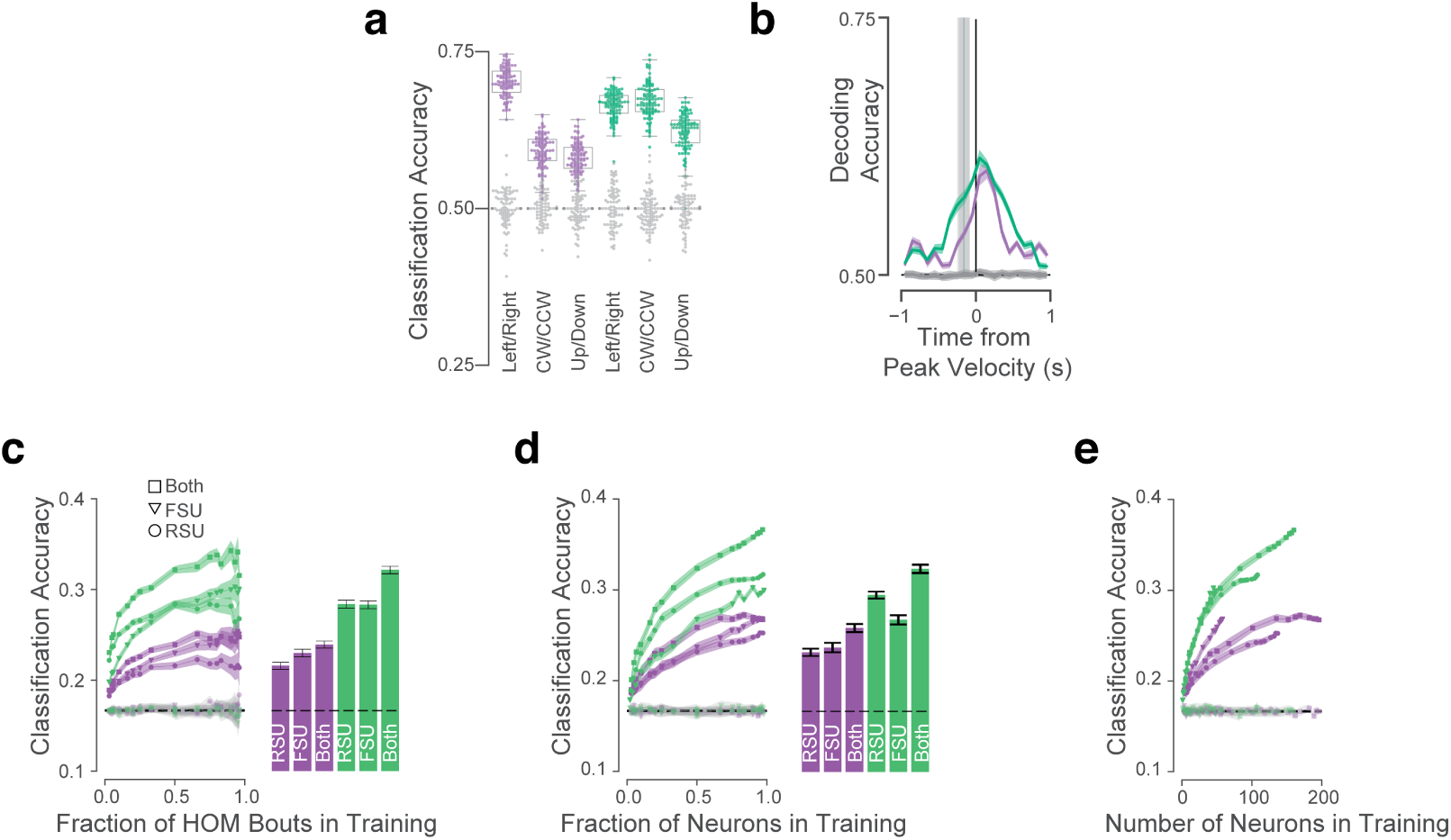
(Related to Figure 2). Decoding Head Orienting Movement Direction from Single-Unit Activity. **(a)** Logistic regression model trained to discriminate Left/Right, CW/CCW, or Up/Down HOMs from single-unit activity (RSU and FSU pooled). Performance was on average somewhat lower in the dark (0.62 ± 0.0035 (mean ± s.e.m.)) than in light (0.65 ± 0.0020) (*p* = 3.2*e* − 12, MWU test), but mixed for individual HOM directions: Left/right (dark: 0.70 ± 0.0023. light: 0.66 ± 0.0022, *p* = 4.1*e* − 21), CW/CCW (dark: 0.59 ± 0.0026. light: 0.67 ± 0.0027. *p* = 4.3*e* − 33), Up/Down (dark: 0.58 ± 0.0024. light: 0.62 ± 0.0029. *p* = 3*e* − 20). Grey: performance of models trained with shuffled labels. Chance performance = 0.5. **(b)** Mean pairwise decoding accuracy as a function of decoding time window (100-ms sliding window) averaged across HOM directions for models trained to discriminate Left and Right, CW and CCW, or Up and Down HOM directions. Green and purple vertical lines indicate mean HOM onset times in light and dark, respectively (grey shading: ±1 std). Gray horizontal lines and shading: performance of decoder trained on shuffled labels. Chance performance = 0.5. **(c)** Left: Classification accuracies for models trained on varying fractions of HOM bouts, from *n* = 100 total bouts, split by neuron types. Right: mean classification accuracies at max bouts. **(d)** Same as **(c)**, but for models trained on varying fractions of neurons. **(e)** Same as **(d)** but plotted as a function of the number of neurons used rather than the fraction.

**Figure S6:**
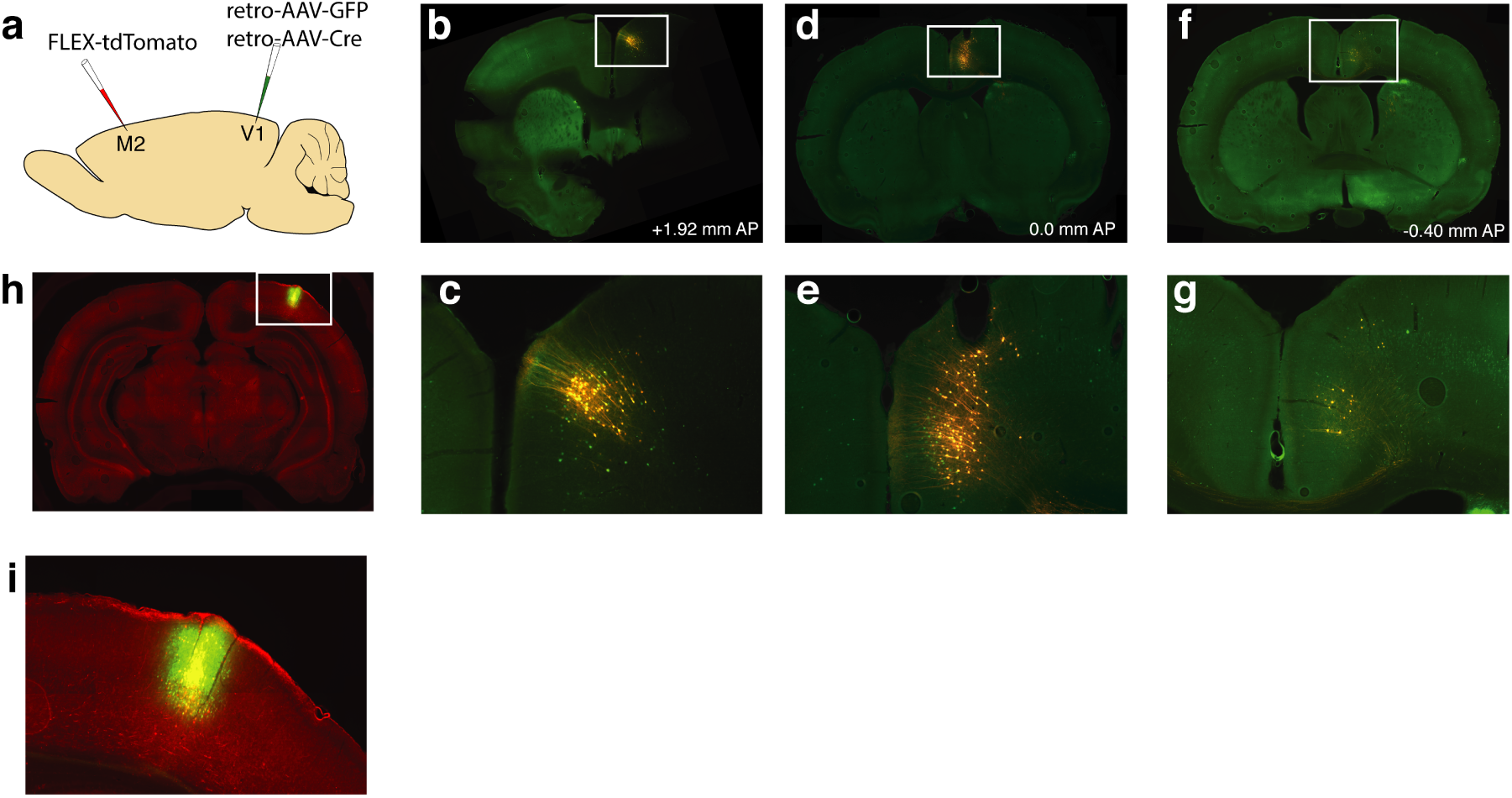
(Related to Figure 4). M2-V1 Viral Tracing. **(a)** Tracing strategy. retro-AAV-GFP and retro-AAV-CRE were co-injected into V1 at the same coordinates as the tetrode implants. FLEX-tdTomato was injected over M2. This intersectional tracing strategy allowed us to visualize projections to V1 originating in diverse cortical and subcortical areas in green and those from M2 in red. **(b**,**c)** Anterior-most V1-projecting M2 neurons around +1.92 mm AP. **(d**,**e)** FLEX-tdTomato injection site in M2 (∼0.0 mm AP). **(f**,**g)** Posterior-most V1-projecting M2 neurons around −0.40 mm AP. **(h**,**i)** Projections within V1 (green) and M2 axons in V1 (red). Panels **c, e, g, i** show detail of white boxes in the other panels.

**Figure S7:**
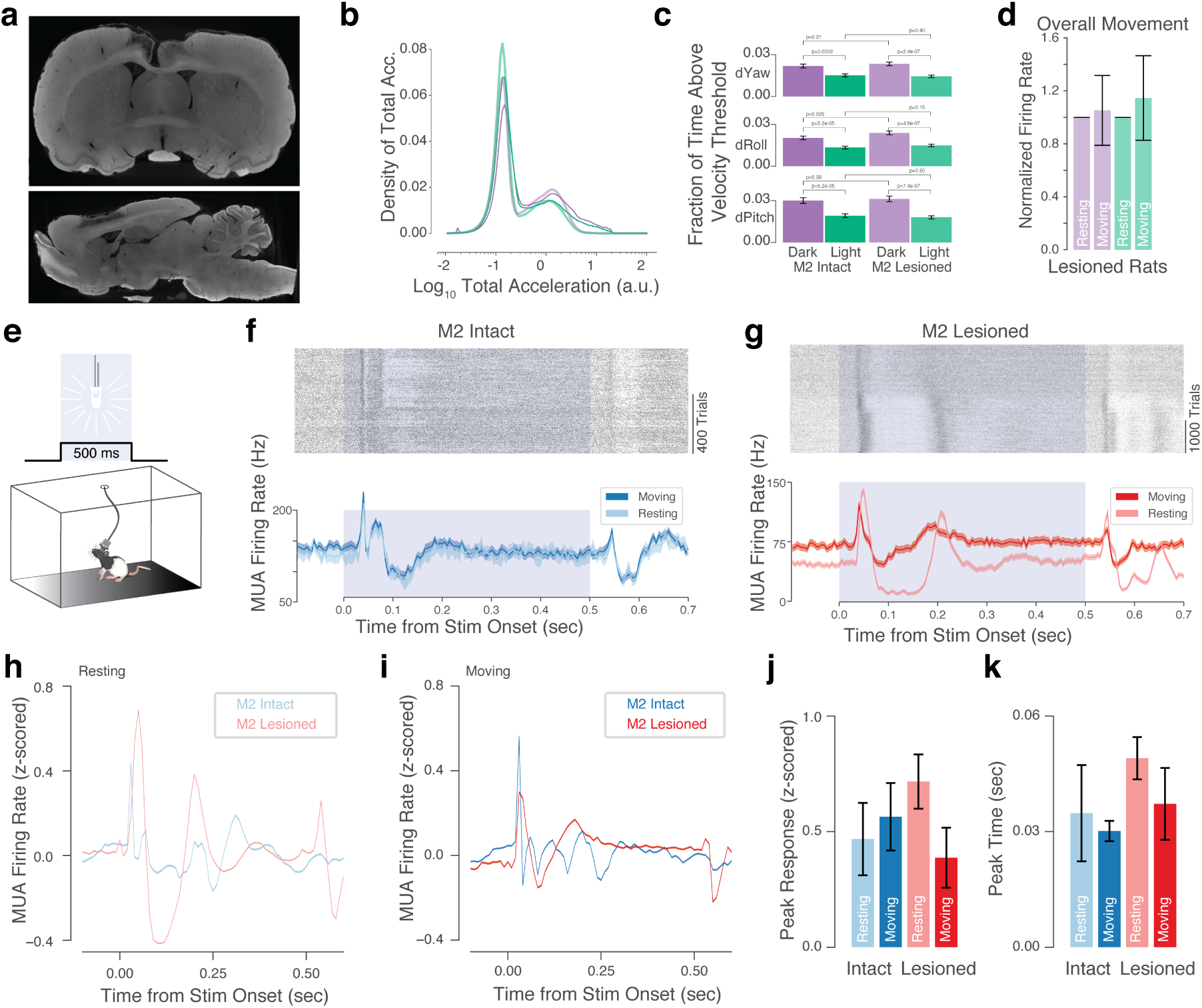
(Related to Figure 4). Movement Statistics and Responses to Visual Stimulation in Lesioned and Non-lesioned Animals. **(a)** Coronal (top) and sagittal (bottom) digital micro-CT slices from one example lesioned brain. **(b)** Histogram of overall movement across sessions in non-lesioned (darker purple/green) and lesioned (lighter purple/green) animals. Lesioned rats spent less time at the highest overall movement vigor levels compared to non-lesioned rats. **(c)** Lesioned and non-lesioned animals spent similar fractions of time per session above angular velocity thresholds (p-values of Wilcoxon rank-sums tests above the bars). **(d)** MUA firing rates in lesioned rats were on average 5.3% and 14.6% higher during movement compared to rest, in the dark and light, respectively (dark: *p* < 2*e* − 4, light: *p* < 1*e* − 10, Wilcoxon test; error bars: median absolute deviation). **(e)** Visual stimuli were delivered while rats behaved freely in their home cages (same as in Fig. 3) to examine visual responses in lesioned animals. **(f**,**g)** Example multiunit responses to flash stimuli. Top: spike raster plots (trials arranged by total acceleration in the 100 ms after stimulus onset). Bottom: mean responses (shading: s.e.m.) split by moving (darker blue/red) and resting (lighter blue/red) in non-lesioned (**f**; blue) and lesioned (**g**; red) animals. **(h**,**i)** Mean responses to flashes during rest **(h)** and overall movement **(i)** in *n* = 8224 trials in *n* = 3 non-lesioned animals across *n* = 13 sessions and *n* = 12975 trials in *n* = 4 lesioned animals across *n* = 11 sessions. **(j)** Peak z-scored responses. Error bars: standard deviation. **(k)** Timing of peak responses post stimulus onset. Error bars: standard deviation.

**Figure S8:**
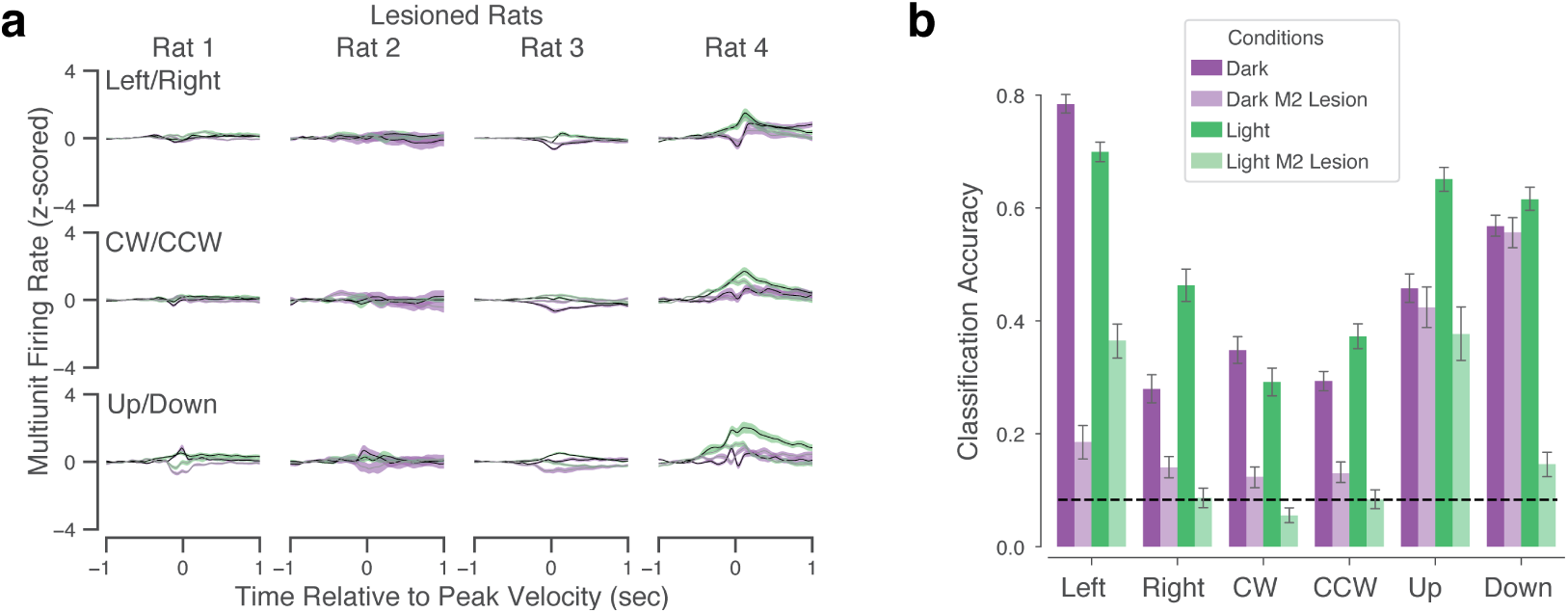
(Related to Figure 4). Head Orienting Movement Activity and Direction Decoding in Lesioned Rats. **(a)** Same as Fig. S2 but for lesioned animals. **(b)** Classification accuracy for each HOM direction in control and lesioned rats. Accuracy was significantly lower for each direction (Mann-Whitney U test, all *p* < 1*e* − 18) except for Up (*p* = 0.059) and Down (*p* = 0.28) in the dark.

